# Benchmarking cell type deconvolution in spatial transcriptomics and application to cancer immunotherapy

**DOI:** 10.64898/2026.01.13.699379

**Authors:** Amanda Sun, Tamjeed Azad, Chrysothemis Brown, Yuri Pritykin

## Abstract

Cell type deconvolution is essential for resolving multicellular composition in spatial transcriptomics data, yet the accuracy and consistency of current methods across biological contexts remain uncertain. We introduce a benchmarking framework using realistic simulations that incorporate spatial and transcriptional complexity. We find that a simple marker gene signature scoring approach performs competitively, often outperforming more complex models, particularly for rare cell types. We further apply this method to a new spatial transcriptomics dataset profiling the early response to a single dose of anti-PD1 checkpoint blockade immunotherapy in a mouse cancer model, revealing pronounced spatially localized immune and microenvironmental changes in both tumors and draining lymph nodes. Our findings have implications for the interpretation of many published spatial transcriptomics studies that rely on cell type deconvolution, and they provide a robust, interpretable strategy for future analyses, even in the absence of matched single-cell references.

**Highlights:**

- Framework for realistic benchmarking of spatial deconvolution methods
- Marker gene scoring performs robustly for rare cell types across diverse datasets
- Single-dose anti-PD1 cancer immunotherapy triggers profound localized response

## Introduction

The spatial organization of cells within tissues is fundamental to their function in development, homeostasis and disease. Recent technological advances have enabled the integration of spatial information with high-throughput gene expression profiling. Among commercially available platforms, 10x Genomics Visium is widely used due to its accessibility and transcriptome-wide coverage. However, the spatial resolution of Visium is limited: each spot captures expression from multiple neighboring cells, necessitating computational deconvolution to resolve the underlying cellular composition^1^.

To address this limitation, numerous cell type deconvolution methods have been proposed for Visium data, often relying on single-cell RNA-seq (scRNA-seq) reference datasets to estimate cell type proportions in each spot. While several benchmarking studies have evaluated these methods ^2–6^, critical questions remain, as existing analyses have not fully addressed the diverse biological contexts and applications in which accurate cell type deconvolution is needed. Notably, most existing evaluations do not account for biologically realistic spatial structure, differences in cell type abundance, and the presence of rare cell populations, factors that are especially important in complex tissues such as tumors, lymphoid organs, and barrier tissues. These environments exhibit intricate spatial organization, with diverse epithelial, immune, and stromal cell types interacting across specialized microenvironments such as germinal centers, epithelial crypts, tertiary lymphoid structures, and tumor-immune interfaces, as well as through broader epithelial-immune and immune-immune crosstalk. Accurate deconvolution in such contexts requires distinguishing closely co-localized cell types and capturing dynamic cell-cell interactions.

In the tumor microenvironment, infiltrating immune cells, often present at low abundance, can drive anti-tumor immunity or mediate immunosuppression^7,8^. Understanding their spatial distribution is essential for studying responses to immunotherapies such as immune checkpoint blockade. Yet, the performance of current deconvolution methods in detecting such rare cell populations remains poorly characterized.

To address these gaps, we developed a spatially and transcriptionally realistic simulation framework for benchmarking cell type deconvolution methods, revealing that a simple marker gene scoring approach performs robustly, especially for rare cell types. We then applied this method to a new spatial transcriptomics dataset profiling the early response to a single dose of anti-PD1 cancer immunotherapy in a mouse model. This analysis uncovered spatially localized immune and microenvironmental changes in both tumors and tumor-draining lymph nodes, demonstrating the method’s utility and highlighting implications for interpreting past and future spatial studies that rely on cell type deconvolution.

## Results

### Cell type deconvolution methods yield variable predictions

**Figure 1.**
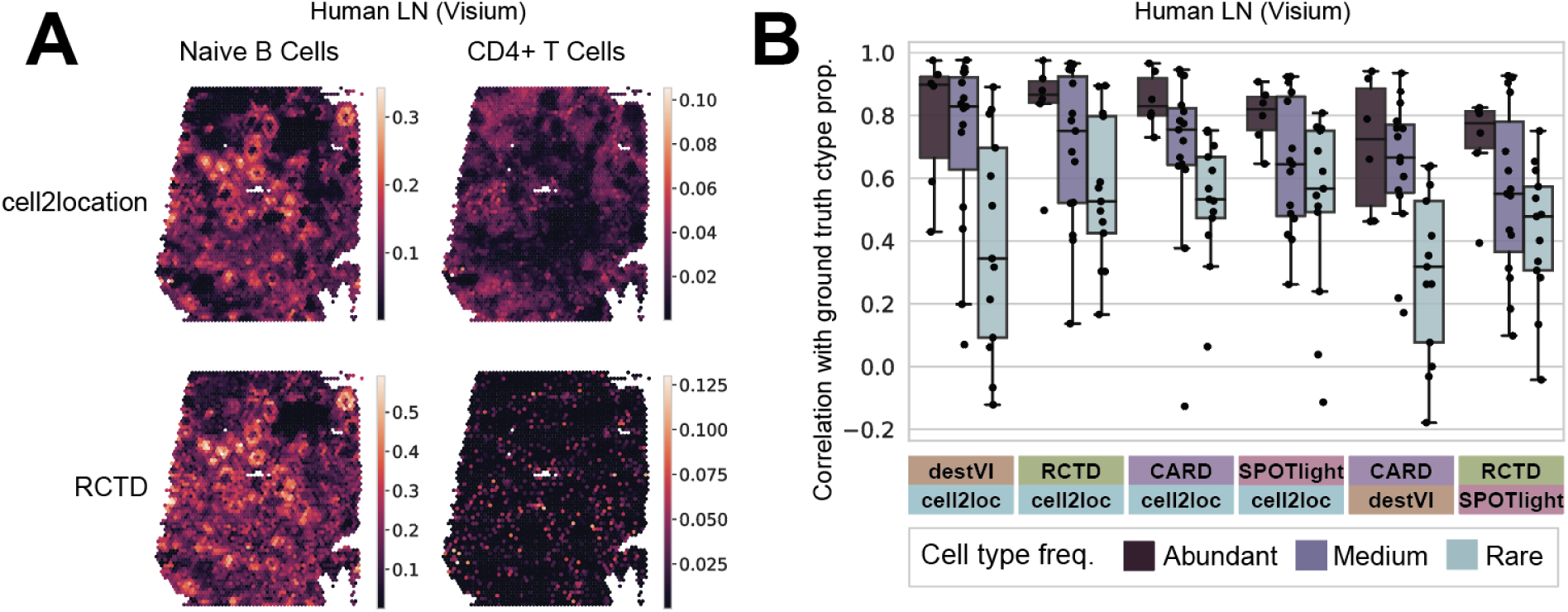
Cell type deconvolution method comparison on human lymph node Visium data. **A.** Spatial plots of a human lymph node Visium dataset showing predicted cell type proportions from cell2location and RCTD. Predictions match well for naive B cells (left) but differ substantially for CD4 T cells (right). **B.** Boxplot of Pearson correlations between predicted cell type proportions from pairs of deconvolution methods applied to the human lymph node dataset. Top six most concordant method pairs are shown, with cell types grouped by abundance category: abundant, medium, rare (see Methods). Center line: median; box: interquartile range (IQR); whiskers: 1.5x IQR.

To assess the performance across methods and further motivate benchmarking, we analyzed a human lymph node Visium dataset^9^ with a matching scRNA-seq reference and annotated cell types^10^. Notably, two widely used deconvolution methods cell2location^10^ and RCTD^11^ showed strong agreement for some cell types, such as naïve B cells, but diverged substantially for others, including CD4+ T cells (**Fig. 1A**). We further tested four additional reference-based methods (CARD^12^, destVI^13^, SPOTlight^14^, UCDSelect^15^) and one reference-free method (UCDBase^15^). Pairwise correlation analysis of the results produced by these methods revealed substantial variation in predicted cell type proportions (**Fig. 1B, Fig. S1**). To investigate this further, for each cell type we used the mean predicted proportion across methods as a proxy of the true abundance. Stratifying the between-method correlation values by cell type abundance revealed that agreement between methods decreases markedly for rare cell types (**Fig. 1B**). These results underscore the need for systematic benchmarking of deconvolution methods, especially in realistic biological contexts where resolving rare cell populations is critical.

### A framework for spatially and transcriptionally realistic benchmarking of cell type deconvolution in spatial transcriptomics

**Figure 2.**
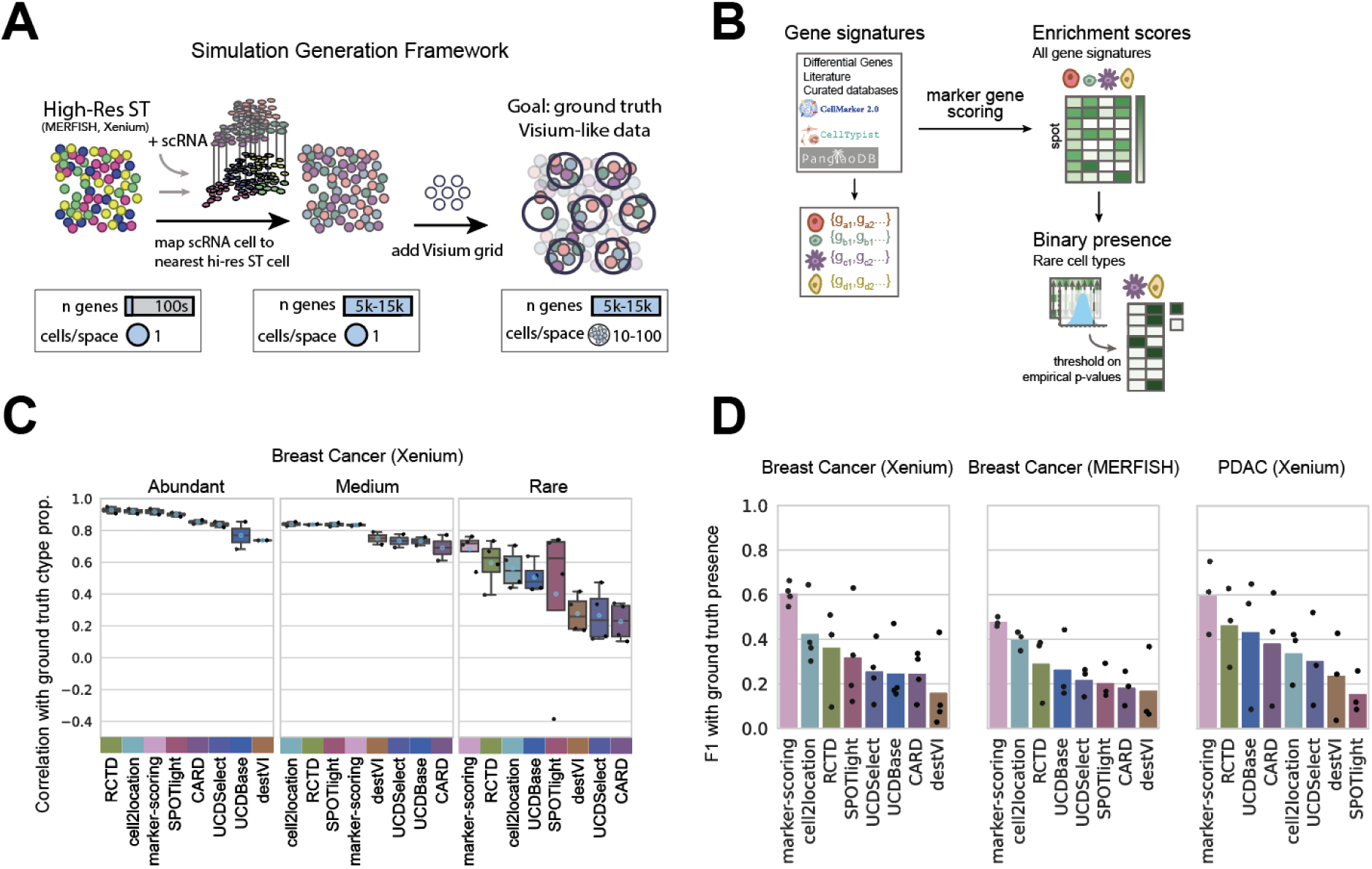
A framework for benchmarking cell type deconvolution methods for Visium spatial transcriptomics. **A.** Schematic of spatially and transcriptionally realistic Visium-like simulations. High-resolution single-cell spatial transcriptomics data with limited gene coverage (e.g. Xenium or MERFISH) is aligned with a matching single-cell genome-wide transcriptomics dataset, to generate simulated Visium-like data with known ground truth cell type compositions. See Methods for more details. **B.** Schematic of marker gene signature enrichment scoring used for cell type deconvolution. **C.** Boxplot of Pearson correlations between predicted scores and ground truth cell type proportions in simulated Visium-like breast cancer data based on Xenium. Blue circle: mean; center line: median. **D.** F1 accuracy scores for rare cell types across deconvolution methods in simulated datasets.

To quantitatively benchmark the performance of cell type deconvolution methods, we developed a framework for generating spatially and transcriptionally realistic Visium-like simulations with known ground truth cell type composition for each spot. In this framework, high-resolution spatial transcriptomics datasets from MERFISH^16,17^ or Xenium (10x Genomics) are aligned to a matching scRNA-seq reference, and nearby cells are aggregated into artificial Visium spots (**Fig 2A, Methods**). This approach addresses a key challenge: high-resolution spatial technologies such as MERFISH and Xenium allow robust single-cell-level cell type annotation, but they measure only a limited number of genes from predefined panels. In contrast, scRNA-seq captures genome-wide gene expression but lacks spatial resolution. By integrating these complementary modalities, we create semi-simulated datasets that combine the spatial precision of MERFISH/Xenium with the transcriptome-wide coverage of scRNA-seq. Aggregating neighboring cells into spots yields Visium-like spatial transcriptomics data with known cell type mixtures in each spot, enabling rigorous, quantitative benchmarking. Using this simulation framework, we generated three Visium-like datasets: a breast cancer dataset based on MERFISH^18^, a second breast cancer dataset based on Xenium^19^, and a pancreatic ductal adenocarcinoma (PDAC) dataset based on Xenium^20^ (**Fig. S2, Methods**).

In addition to published cell type deconvolution methods, we also tested a simple and flexible baseline deconvolution approach through per-spot marker gene signature enrichment using the default gene set scoring function implemented in Scanpy^21^, a widely used single-cell genomics data analysis package (**Fig. 2B**). Unlike reference-based deconvolution methods, this approach does not require a matched scRNA-seq reference and can leverage cell type markers from a variety of sources, including expert knowledge, literature, non-matching single-cell datasets, or public databases such as CellMarker^22^, CellTypist^23^, and PanglaoDB^24^. Importantly, the gene signature scoring method treats each cell type independently and does not require a complete decomposition of the gene expression profile for each spot, allowing for selective scoring of only a subset of relevant cell types. This flexibility is particularly valuable when suitable reference datasets are unavailable, incomplete, or poorly matched to the tissue under study.

We benchmarked performance by computing Pearson correlations between the ground truth cell type proportions and the predicted proportions or marker gene enrichment scores. To evaluate performance across cell type abundance levels, we stratified cell types into three categories: abundant, medium, or rare (**Fig. S3**). In simulations based on Xenium breast cancer data, all methods performed well for abundant cell types (median correlations 0.74–0.93; **Fig. 2C**). However, performance declined for rare cell types (median correlations 0.23–0.72), where marker gene scoring achieved the highest median correlation 0.72, followed by RCTD achieving 0.63. Similar trends were observed in simulations based on MERFISH breast cancer and Xenium PDAC data (**Fig. S4**), with marker gene scoring consistently among the top-performing methods. Cell2location and UCDBase ranked third or fourth across all datasets. For rare cell types, interpreting enrichment scores in terms of binary presence or absence may be more informative. To this end, we binarized marker gene enrichment scores using empirical *p*-values from bootstrapping (**Methods**). For other methods, which return continuous cell type proportions, we applied a fixed threshold of 5% to define presence. Marker gene scoring achieved the highest median F1 score for rare cell types (**Fig. 2D**), with cell2location ranking second in the breast cancer datasets and RCTD second in PDAC. Evaluation using additional metrics, including accuracy, precision, recall, Jaccard index, ROC AUC, and PR AUC, supported these findings (**Fig. S5**). Performance was robust across a range of thresholding parameters (**Fig. S6**). Together, these results highlight the surprisingly strong performance of the simple marker gene scoring approach, especially for rare cell types. Despite its simplicity and lack of reliance on matched scRNA-seq references, it performs comparably to or better than more complex deconvolution methods across a range of evaluation metrics and datasets.

In the main benchmarking tasks above, marker genes were derived from differential expression in the matched scRNA-seq reference (**Fig. S7**). To evaluate performance in scenarios where such a reference is unavailable, we next tested marker gene sets obtained from public databases. While overall accuracy decreased, marker gene scoring remained competitive, with PanglaoDB markers performing especially well (**Fig. S8**). Next, we further explored a key advantage of marker gene scoring, its ability to evaluate each cell type independently, without requiring complete deconvolution across all cell types. To assess this in the context of an incomplete scRNA-seq reference that does not cover all cell types present in the spatial data, we restricted the reference to immune cells only. As expected, performance dropped across all methods; however, marker gene scoring, whether using recalculated differentially expressed genes or database-derived marker sets, remained among the top performers (**Fig. S9**). Finally, we found that among the normalization strategies tested, scTransform yielded the most consistent and robust performance in Visium-like simulations (**Fig. S10**). Together, these results demonstrate that marker gene signature scoring achieves competitive performance even when using publicly available gene sets, and offers additional flexibility by allowing selective scoring of only a subset of relevant cell types.

### Spatially localized responses to a single-dose checkpoint blockade immunotherapy in a mouse cancer model

After demonstrating the competitive performance of marker gene scoring in simulated benchmarks, we applied it to study the spatial immune response to anti-PD1 checkpoint blockade immunotherapy in a mouse cancer model. While immune checkpoint inhibitors such as anti-PD1 have shown remarkable success in subsets of patients across multiple cancer types, many individuals do not respond, and the mechanisms underlying response, whether direct or indirect, remain incompletely understood^25,26^. Moreover, optimal dosing remains unclear, and excessive or prolonged exposure to checkpoint blockade can lead to severe autoimmune side effects^27–29^. To investigate the immediate spatial effects of treatment while minimizing potential toxicity, we administered a single dose of anti-PD1 blocking antibody or isotype control seven days after subcutaneous injection of MC38 tumor cells in C57Bl/6 mice, and harvested both tumors and tumor-draining inguinal lymph nodes (tdLNs) on day 10 for spatial transcriptomic profiling using Visium (**Fig. 3A**).

**Figure 3.**
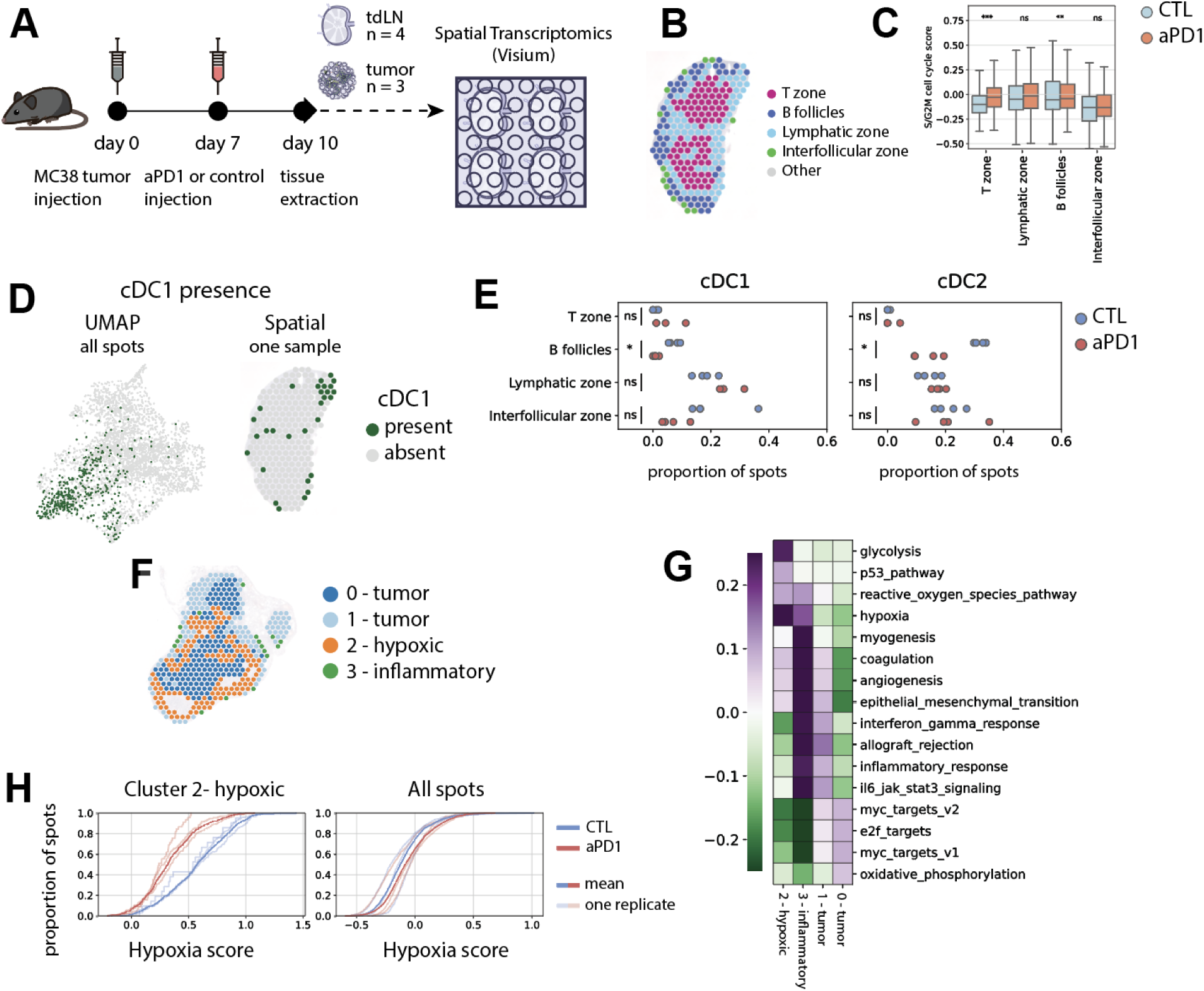
Application of marker gene enrichment-based deconvolution to the tumor immune microenvironments in cancer immunotherapy. **A.** Schematic overview of the experimental design. Upon tumor initiation and subsequent treatment with a single-dose anti-PD1 checkpoint blockade or control, tumors and tumor-draining lymph nodes were isolated for Visium spatial transcriptomic profiling. For each tissue, *n* is the number of samples per condition. From each sample, two adjacent tissue sections were profiled as technical replicates. **B.** Spatial plot of a representative lymph node with annotated tissue regions. **C.** Boxplot of cell cycle enrichment scores per cluster in lymph nodes, colored by treatment condition. Center line: median. Asterisks indicate statistical significance from a two-sided *t*-test between conditions: p < 0.05 (*), p < 0.01 (**), p < 0.001 (***). **D.** Uniform manifold approximation and projection (UMAP) plot (left) and a spatial plot of a representative lymph node (right) showing predicted presence of conventional type 1 dendritic cells (cDC1). **E.** Comparison of the proportion of spots in each lymph node transcriptomic cluster predicted to contain DCs of type 1 and 2. Dots indicate samples from different individual mice. Asterisks indicate statistical significance from a two-sided *t*-test between conditions: p < 0.05 (*), p < 0.01 (**), p < 0.001 (***). **F.** Spatial plot of a representative tumor annotated by Leiden clusters of transcriptomic profiles. **G.** Heatmap of pathway enrichment scores across tumor clusters. Shown pathways are among the top four most enriched per cluster. **H.** Empirical cumulative distribution function (eCDF) plots of hypoxia scores in tumor cluster 2 spots (left) and all tumor spots (right). A separate distribution curve is shown for a tumor section from each individual mouse; a thicker curve shows the distributions in all spots per condition.

In the tdLNs, T cell and B cell zones were robustly identified across multiple Visium sections using marker enrichment scoring (**Methods**). Leiden clustering of the remaining spots further delineated interfollicular and lymphatic zones, yielding reproducible spatial annotations that were consistently observed across biological replicates (**Fig. 3B, Fig. S11A,B**). Following anti-PD1 treatment, we observed an increase in cell cycle gene expression specifically in the T cell zone, consistent with enhanced T cell proliferation in response to checkpoint blockade^30^ (**Fig. 3C, Fig. S11C**). To identify additional immune cell types, we scored differentially expressed genes from LN-resident cell types^31^ in our spatial data (**Fig. 3D,E**). This deconvolution revealed spatially specific treatment effects, including a significant depletion of dendritic cells localized to the B cell follicles following anti-PD1 treatment (**Fig. S11D**). Thus simple marker gene scoring revealed robust and spatially localized immune responses to anti-PD1 treatment in the tdLNs.

We next examined treatment response in the tumor microenvironment by scoring immune cell signatures derived from scRNA-seq of MC38 tumors^32^. While this dataset profiled only immune cells, it was sufficient for robust marker gene scoring-based deconvolution. Spatial transcriptomic data from the tumor was clustered (**Fig. 3F, Fig. S12A,B**), and enrichment scores were computed for cell cycle genes and hallmark pathways from MSigDB^33,34^ (**Fig. 3G, Fig. S12C**). This analysis identified four tumor clusters with distinct spatial localization and gene expression programs: cluster 0 capturing interior tumor regions; another tumor cluster 1; cluster 2 enriched in glycolysis and hypoxia pathways; and cluster 3 capturing inflammatory regions enriched in immune signatures. Cell type markers from PanglaoDB produced similar deconvolution results to the scRNA-derived signatures (**Fig. S12D**), reinforcing robustness. Comparing pathway enrichment across clusters and treatment conditions revealed that hypoxia scores were significantly reduced after anti-PD1 treatment, specifically in the hypoxic cluster 2 (**Fig. 3H, Fig. S13A,B**). These hypoxic regions were enriched for a neutrophil state previously identified as pro-angiogenic and associated with hypoxic environments^35^ (**Fig. S13C**). Marker gene scoring further revealed that anti-PD1 treatment decreased the abundance of this pro-angiogenic neutrophil subset, while increasing the representation of alternative neutrophil states in the same region, suggesting a possible cellular mechanism underlying the reduced hypoxic scores (**Fig. S13D**). Together, these results demonstrate that gene signature scoring can sensitively detect spatially localized immunotherapy responses in both tumors and tdLNs, even after a single dose of anti-PD1 checkpoint inhibitor, highlighting its utility for dissecting early cellular and microenvironmental changes following treatment.

## Discussion

We systematically benchmarked cell type deconvolution methods for Visium spatial transcriptomics using spatially and transcriptionally realistic simulations, and identified cell type abundance as a major factor influencing performance. Surprisingly, we found that marker gene scoring, a simple and reference-light approach, performs competitively with more complex methods, especially for rare or underrepresented cell types. Importantly, this approach enables flexible analyses in the absence of matched scRNA-seq data and allows selective scoring of only relevant cell types, making it particularly suitable for applications in clinical or poorly characterized tissues.

Our application to anti-PD1 immunotherapy reveals early and spatially restricted immune responses in both tumors and tumor-draining lymph nodes. A single treatment dose altered dendritic cell distribution and hypoxic niches, highlighting the utility of spatial transcriptomics to detect immediate, localized effects of checkpoint blockade. These findings demonstrate the power of marker gene scoring for dissecting complex tissue responses, with implications for immunotherapy, infectious disease, and beyond.

## Resource availability

Spatial transcriptomics data for the tumor and tumor-draining lymph node have been uploaded to GEO at the accession code GSE293019 and will be made available upon journal publication. Computational code used for the analysis in this study is available at https://github.com/pritykinlab/Spatial-Cell-Type-Deconvolution-Benchmarking.

## Acknowledgments

We thank all members of the Pritykin lab for helpful discussions. This work was supported by the NIH/NIAID grant DP2AI171161, Ludwig Institute for Cancer Research, Weill Cancer Hub East and AACR-Bristol-Myers Squibb Immuno-oncology Research Fellowship.

## Methods

### Preprocessing single-cell spatially resolved transcriptomics

MERFISH and Xenium spatial data were used to generate simulated Visium-like data. Spatial data was first filtered to remove cells with total counts in the lowest or highest 2nd percentile. Cells were then normalized by volume (MERFISH) or area (Xenium), then by library size, and then z-score normalized across cells before dimension reduction with PCA, KNN-graph construction, and leiden clustering. Clusters were manually annotated by expression of cell type-specific markers. Cells expressing markers from multiple cell types–likely to be mis-segmented–were removed. The same normalization was used for downstream alignment with scRNA-seq data.

For the PDAC Xenium data, public annotations from 10X Genomics were used as initial annotations. Annotations for T cell subtypes were refined using the method described above.

### Preprocessing scRNA-seq data for simulation generation and deconvolution reference

For simulation generation, both spatial breast cancer datasets were aligned to scRNA-seq from Pal et al^36^, and the PDAC dataset was aligned with scRNA-seq from Werba et al^37^. Low quality information was filtered out: genes expressed in few cells, cells expressing few genes, and cells with high counts coming from mitochondrial genes were filtered out. Cell type labels from the original publications were used if available, and were subset to match the cell type labels present in the spatial data. Otherwise for manual cell type annotation, data was normalized by library size and then log-transformed. scVI^38^ was used for batch correction and dimensionality reduction. From these results, a KNN-graph was constructed and leiden clusters were found and manually labeled with cell type identity using marker genes. For all datasets, after cell type annotation, raw data was normalized by library size, log-transformed, and then scaled to unit variance and zero mean for alignment with spatial data.

For use as reference for cell type deconvolution, scRNA-seq was taken for both breast^39^ cancer datasets from Wu et al^40^ and for PDAC from Zhang et al^40^. Dataset filtering and cell type annotation were performed as described above. Differential gene expression was performed to get cell type-specific genes for marker gene-based deconvolution. Log fold change values were calculated between the mean expression of library size-normalized counts for cells included and excluded from each cell type. P values were calculated using a Mann-Whitney U test and corrected for multiple hypothesis testing with Benjamini-Hochberg procedure. For each cell type, differential genes were filtered out with a 0.5 log fold change value cutoff and 0.01 adjusted p value cutoff. Then genes in the top 15th percentile of remaining log fold change values were taken as the marker genes for that cell type.

For all scRNA-seq datasets, cells were subset to cell types shared in the spatial data. Spatial datasets were also subset to broad cell types shared in the scRNA data. For both PDAC datasets, data was first analyzed from both liver metastasis and pancreas for preprocessing and cell type identification. Data was then subset to only pancreatic samples for more consistency with the spatial Xenium sample.

### Simulation Framework

Single-cell resolution spatially resolved transcriptomics data was aligned with a scRNA-seq dataset using Harmony^41^. Normalized single-cell RNA and spatial datasets were concatenated, PCA was performed, and the top 100 PCs were used for alignment with Harmony. Each spatial cell was aligned randomly with one of the five closest scRNA-seq cells, and this process was repeated independently five times. The cell type label was taken from the scRNA-seq cell. From the spatial coordinates, cells were assigned to corresponding 55µm diameter spots in a Visium-like spatial grid. Four sets of translated grids were used for a total of 20 simulations. Within each spot, the scRNA-seq gene expression of every cell was summed. To match the count distribution of Visium data, total counts per spot for the entire simulation were downsampled to a Gaussian distribution with mean 20,000 counts while keeping each spot’s relative ranking by total counts. The simulation was then filtered to remove genes and spots with low expression and normalized by either scTransform^42^, library size, or not normalized. Results are reported as the mean across the 20 simulated datasets for each spatial and scRNA-seq pairing.

### Differential gene expression analysis for assessing alignment quality

For all cells aligned to each cell type, gene expression was taken from either a cell in the scRNA-seq data or from a cell in the single-cell spatial that were matched to each other.

Differential gene expression was performed as described above to compare each cell type versus all others using either of the gene expression types. Then spearman correlation was calculated between the logFC values for all genes between the scRNA-seq and single-cell spatial differential gene analyses.

### Preprocessing human LN data

Human LN Visium data was downloaded from 10x Genomics^9^. A single-cell reference was curated and annotated in Kleshchevnikov et al^10^. Both Visium and scRNA-seq datasets were filtered by removing cells expressing fewer than 100 genes and genes expressed in fewer than 10 cells. Differential genes between cell types in the scRNA-seq reference were found as described above. The Visium data was normalized with scTransform before applying deconvolution methods.

### Deconvolution method benchmarking

For all methods tested: cell2location, RCTD, CARD, destVI, SPOTlight, UCDBase, and UCDSelect, they were run using default parameters and following their respective tutorials unless otherwise stated below.

Cell2location was run with N_cells_per_location of 20, detection_alpha of 200.

RCTD was run with doublet_mode = ‘full’. The resulting cell type weights were normalized to proportions by transforming them to sum to 1 for each spot.

CARD was run with 100 minimum counts per gene and 5 minimum counts for each spot for createCARDOBject().

destVI was run with 2000 epochs for training the deconvolution model on the spatial data.

For SPOTlight, marker genes from the scRNA-seq reference were taken following the tutorial. The top HVGs were found (3000 for MERFISH breast cancer and 8000 for the Xenium simulations), and the scran function scoreMarkers was used to identify cell type-specific genes.

From these results, genes with a mean AUC greater than a threshold (0.8 for MERFISH breast cancer and 0.6 for the Xenium simulations) were kept. Cells in the reference were also downsampled to at most 100 per cell type.

UniCell Deconvolve was run in two modes: UCDSelect which uses a single-cell reference and UCDBase which is reference-free. UCDselect was run with default parameters. For UCDBase, cell types were subset to match the cell types in the scRNA-seq reference.

### Thresholding marker gene scoring for predicting absolute presence

From the normalized spatial data counts, values for genes were sampled independently with replacement via marginal bootstrapping to generate expression of a bootstrapped cell. Values for 10,000 bootstrapped cells’ gene expression were sampled in this way. Marker gene scoring was performed on these bootstrapped cells to generate a background distribution. Empirical p-values were calculated by the proportion of scores in the background distribution greater than or equal to the mean marker gene enrichment score of the observed data.

### Evaluating prediction of ground truth cell type presence

Marker gene scoring enrichment scores were thresholded as described above to evaluate the prediction of per-spot present. An empirical p-value cutoff of 0.05 was used. For all other cell type deconvolution methods, the predicted proportion was thresholded at 5%. For all metrics, all spots which contained at least one of a cell type in the ground truth were considered as positive for that cell type. For ROC AUC and PR AUC calculations, either the inverse p-value (for marker gene scoring) or the predicted proportions (for all other methods) were used as the predicted values.

### Immune-only

A situation where the scRNA-seq reference for deconvolution is not perfectly matching was simulated by subsetting only the scRNA-seq references to immune cell types. Differential genes were recalculated on the subset reference for marker gene scoring-based deconvolution. The other scRNA-seq-reference based deconvolution methods were rerun with the subset reference. UCDBase results were subset to select only the cell types matching the subset reference.

### Public database genes

For public cell type marker gene sources, gene signatures were manually curated to select cell types matching those in the single-cell reference. When multiple cell types in the database corresponded to one in the reference, their marker genes were concatenated together.

### Mouse anti-PD1 cancer experiment

MC38 tumor cell line was implanted subcutaneously (1 million cells). After 7 days, mice with a visible tumor were injected with aPD1 (Bioxcell) or isotype control 250mcg/mouse. 3 days later the tumors and tumor draining lymph nodes were harvested.

### Preprocessing MC38 spatial transcriptomics

Mouse tumor and tumor-draining lymph node (tdLN) Visium data were processed separately. For both datasets, cells with low counts, few expressed genes, and high percent of counts (>10%) from mitochondrial genes were removed from each sample. Genes with low counts and expressed in few cells were also filtered out. Cutoffs for cell and gene filtering were chosen for each sample separately.

For the tumor data, batch correction and dimension reduction were performed with scVI as above for scRNA-seq for alignment. From the scVI latent dimensions, a KNN-graph was constructed and leiden clustering was performed to identify four transcriptomic clusters. Raw data was also normalized with scTransform for each sample for use with marker gene scoring.

For the tdLN data, batch correction was not needed, so data was normalized together by scTransform. Normalized data was then used for PCA (100 components) and KNN-graph construction (30 neighbors and 30 PCs). After the T cell and B cell zones were identified through marker gene scoring, leiden clustering was applied to the remaining spots to identify three clusters.

### tdLN zone annotation

Spots within the T cell and B cell zone areas were identified as those enriched with the corresponding lymphocytes. The remaining spots were clustered and annotated as the lymphatic and interfollicular zone based on spatial patterning. One cluster was specific to only one lymph node, so it was labeled as ‘other’ and removed for downstream analysis.

### Marker gene processing for MC38 tumor and lymph node

Differential gene expression analysis for lymph node cell types were taken directly from Huang et al^31^. This study performed scRNA-seq on healthy, non-tumor murine, inguinal LNs and identified LN-resident cell types. The top 100 genes per cell type with highest log fold change values and FDR-corrected p-values < 0.05 were reported. From these results, similar cell types were merged (e.g. Neutrophils 1 and Neutrophils 2), and genes were subset to only those in the top 15th percentile of log fold change values for each cell type.

scRNA-seq data profiled in MC38 tumors and cell type annotations were taken from Zhang et al^32^. Only data from tumors treated with the isotype control were used. The count matrices were filtered to only keep cells with counts between 100-50,000, genes between 600-6500, and percent counts from mitochondrial genes less than 10%. Genes were removed if they were expressed in fewer than 50 cells or had fewer than 10 counts. Differential expression between the cell types was performed as described above. Genes for each cell type were first subset to those with adjusted p values less than 0.01 and log fold change values greater than 0.5. Of these remaining genes, genes in the top 15th percentile of log fold change values for each cell type were used as cell type-specific marker genes for deconvolution.

## Supplementary figures

**Figure S1.**
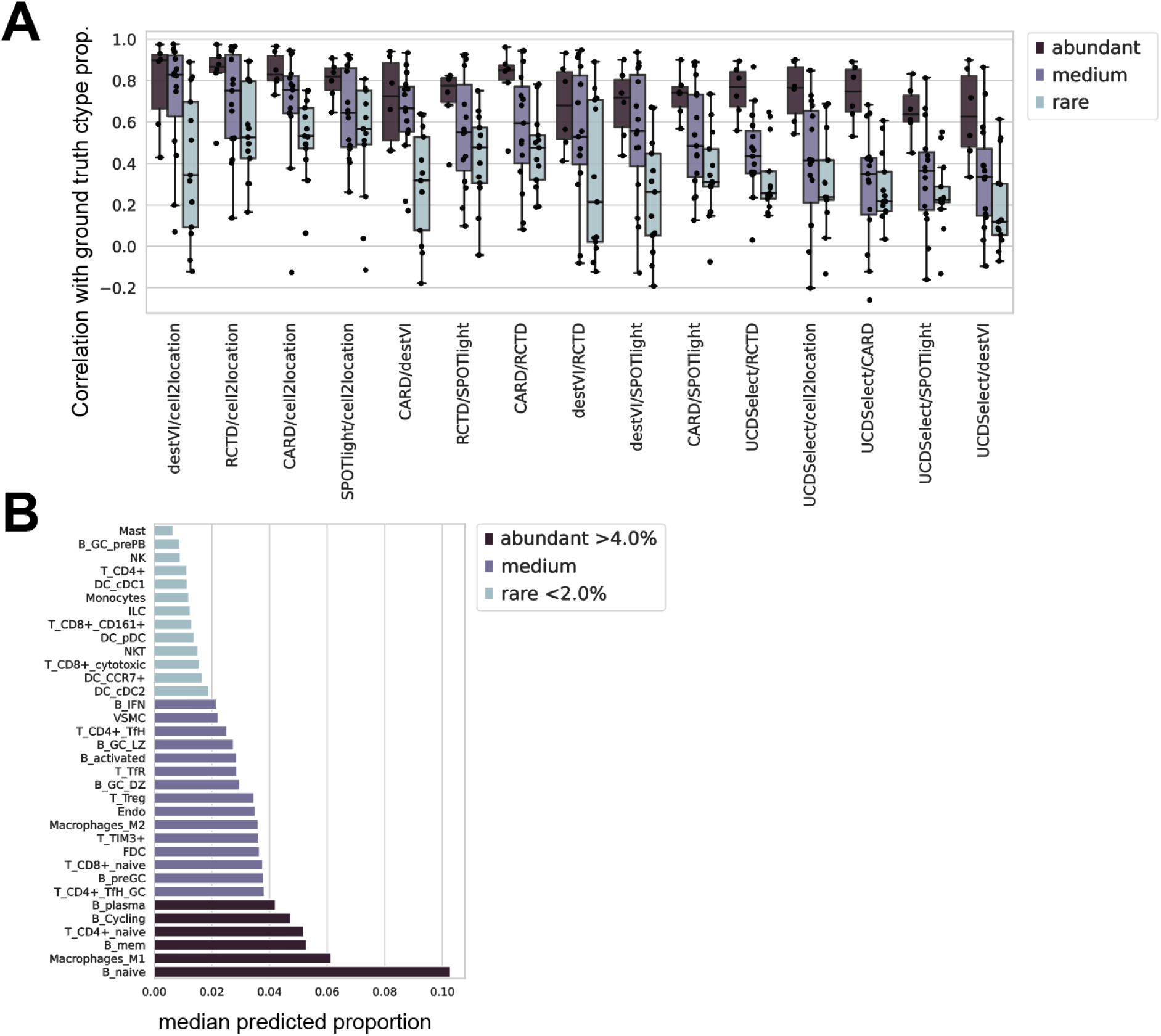
Application of cell type deconvolution methods on the human lymph node Visium dataset. A. Boxplot of Pearson correlations between predicted cell type proportions for all pairs of tested deconvolution methods. Cell types are grouped by their mean predicted abundance category: abundant, medium, or rare. Center line: median, box: interquartile range (IQR), whiskers: 1.5x IQR. B. Barplot of median predicted cell type frequencies across all methods. For each cell type, the median was calculated from the mean predicted proportion across spatial transcriptomics spots. Bars are colored by their assigned abundance category.

**Figure S2.**
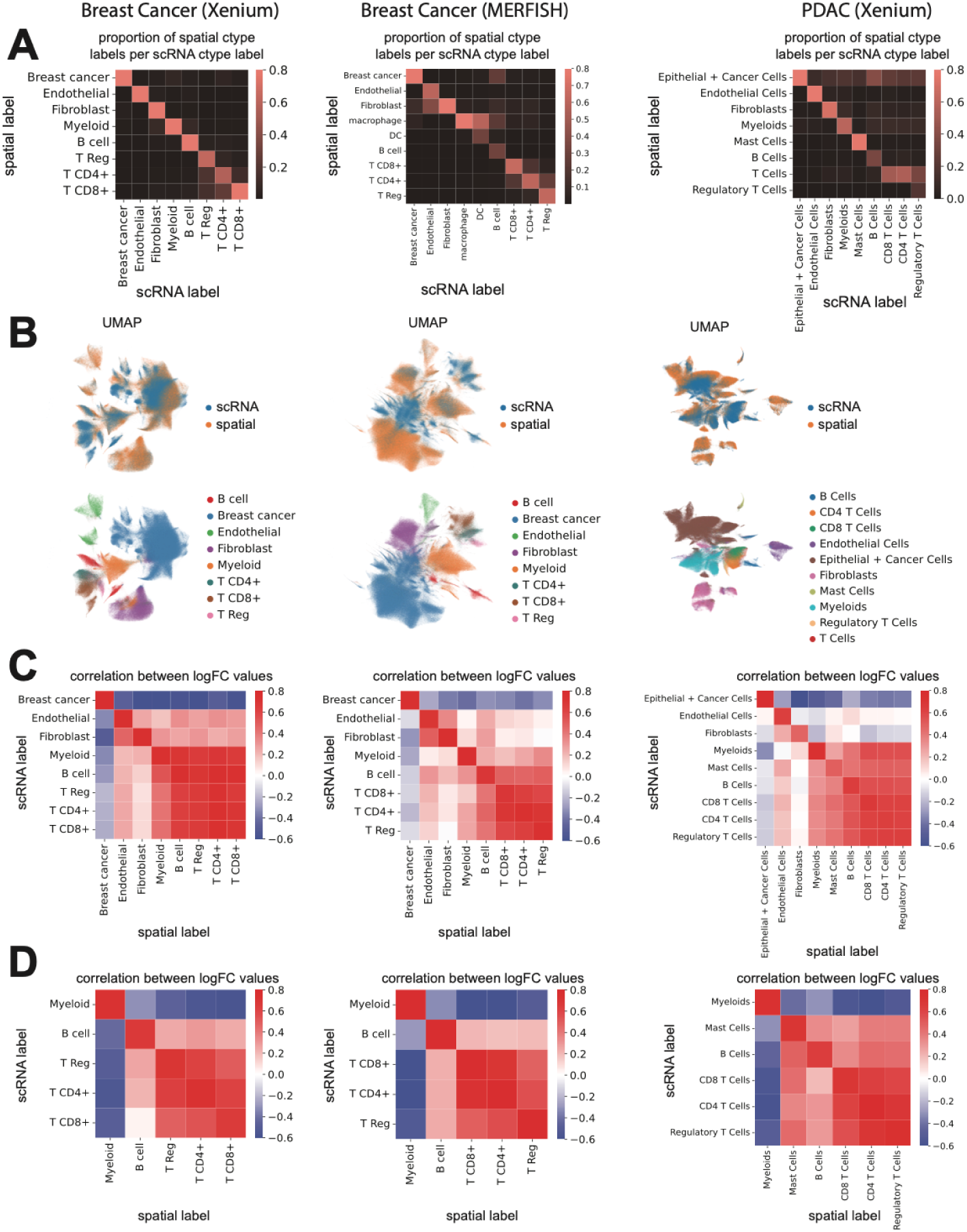
Quality of alignment between MERFISH or Xenium spatial transcriptomics and matched scRNA-seq. A. Heatmaps showing the proportion of spatial cells (from MERFISH or Xenium) annotated to each scRNA-seq cell type label, for each dataset. B. UMAP plots constructed from joint latent dimensions of aligned spatial and scRNA-seq data, colored by data source (top) and by cell type (bottom). Cell type labels are obtained from scRNA-seq analysis and then mapped to spatial data (**Methods**). C. Heatmaps showing correlations between differential expression (log fold change) values for each cell type versus all other cells in the spatial data. Comparisons are shown for cell type labels derived from direct spatial annotations and from mapping via alignment with scRNA-seq. D. Same as panel C, but restricted to immune cells in the spatial dataset.

**Figure S3.**
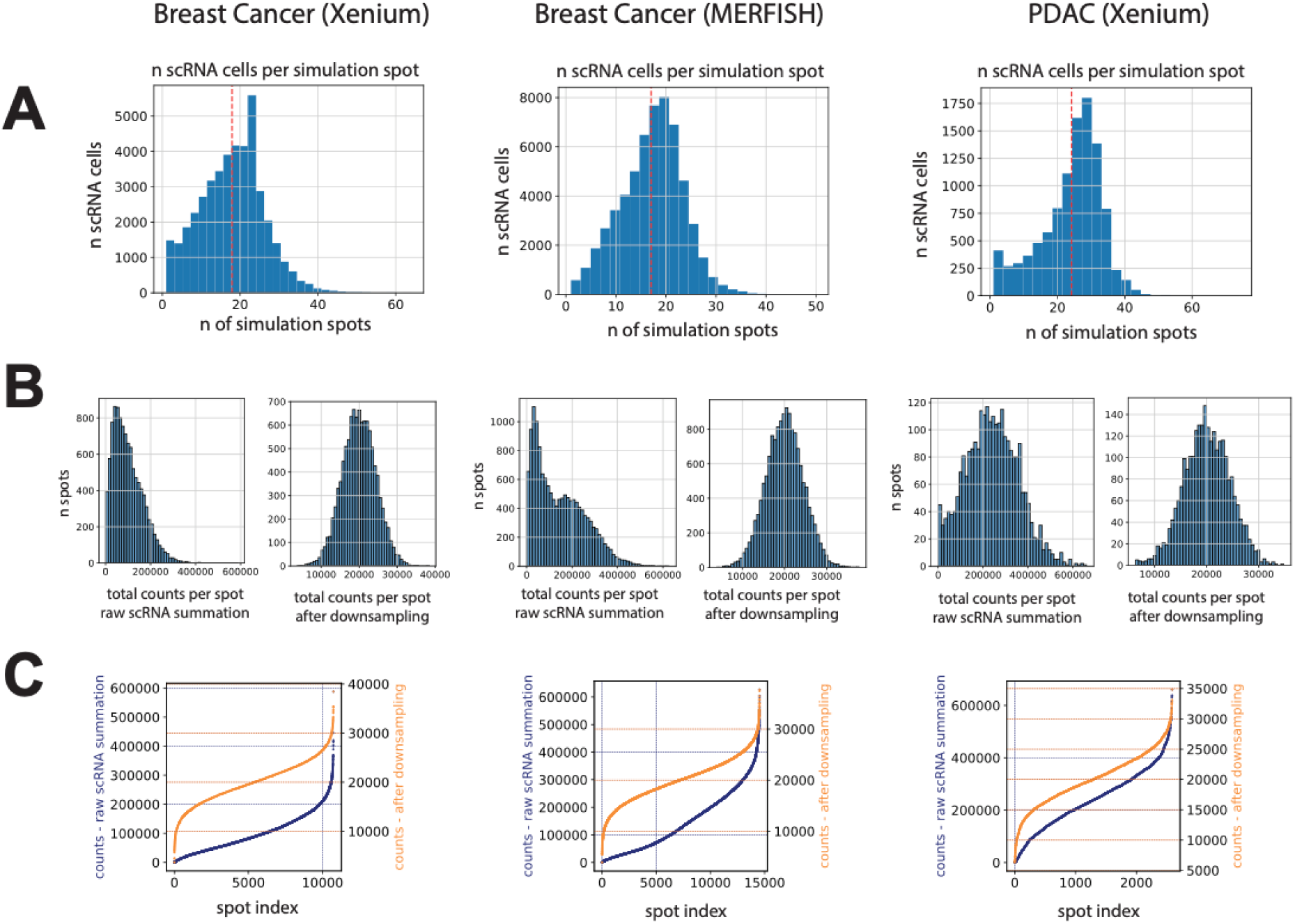
Count distributions in simulated Visium-like data. A. Histogram showing the number of single cells aggregated into each simulated Visium spot for each dataset. B. Histograms of total counts per spot before (left) and after (right) downsampling for each dataset. Downsampling was performed to match a Gaussian distribution approximating total counts while preserving the relative ranking of spots by total count. C. Cumulative distribution plots showing total counts per spot before (blue) and after (orange) downsampling, for all spots across datasets.

**Figure S4.**
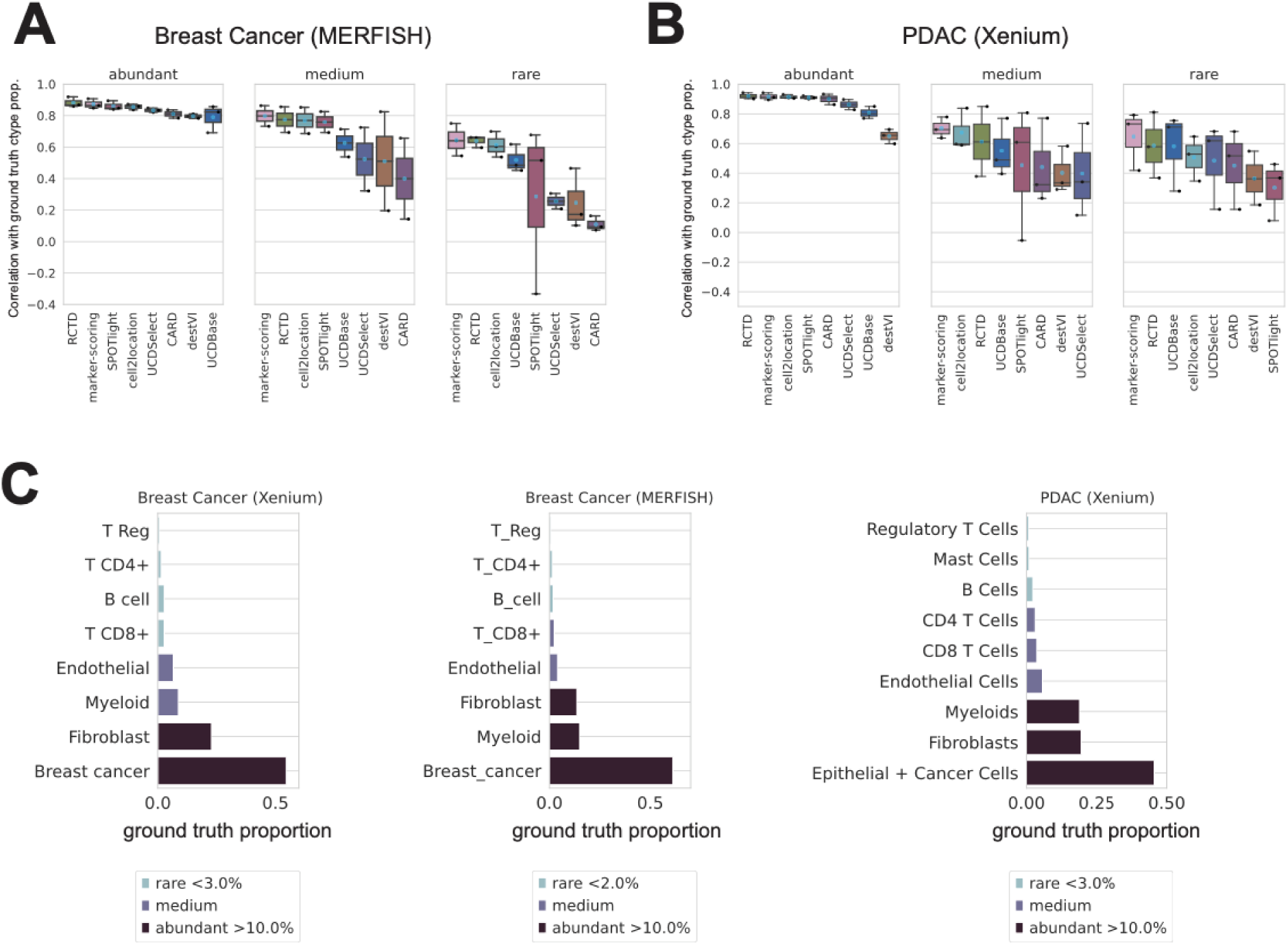
Benchmarking correlations between predicted and ground truth cell type proportions across abundance categories. A. Boxplots showing Pearson correlations between predicted cell type scores from each deconvolution method and the ground truth cell type proportions in the MERFISH breast cancer simulation. Boxes are grouped by cell type abundance category (abundant, medium, rare), with individual cell types overlaid as points. Blue diamond: mean; center line: median; box: interquartile range (IQR); whiskers: 1.5x IQR. B. Same as panel A, but for the Xenium PDAC simulation. C. Barplots showing the ground truth relative abundance of each cell type in the simulated datasets. Relative frequency was calculated as the proportion of cells of each type among all single cells in the simulation. Bars are colored by their assigned abundance category.

**Figure S5.**
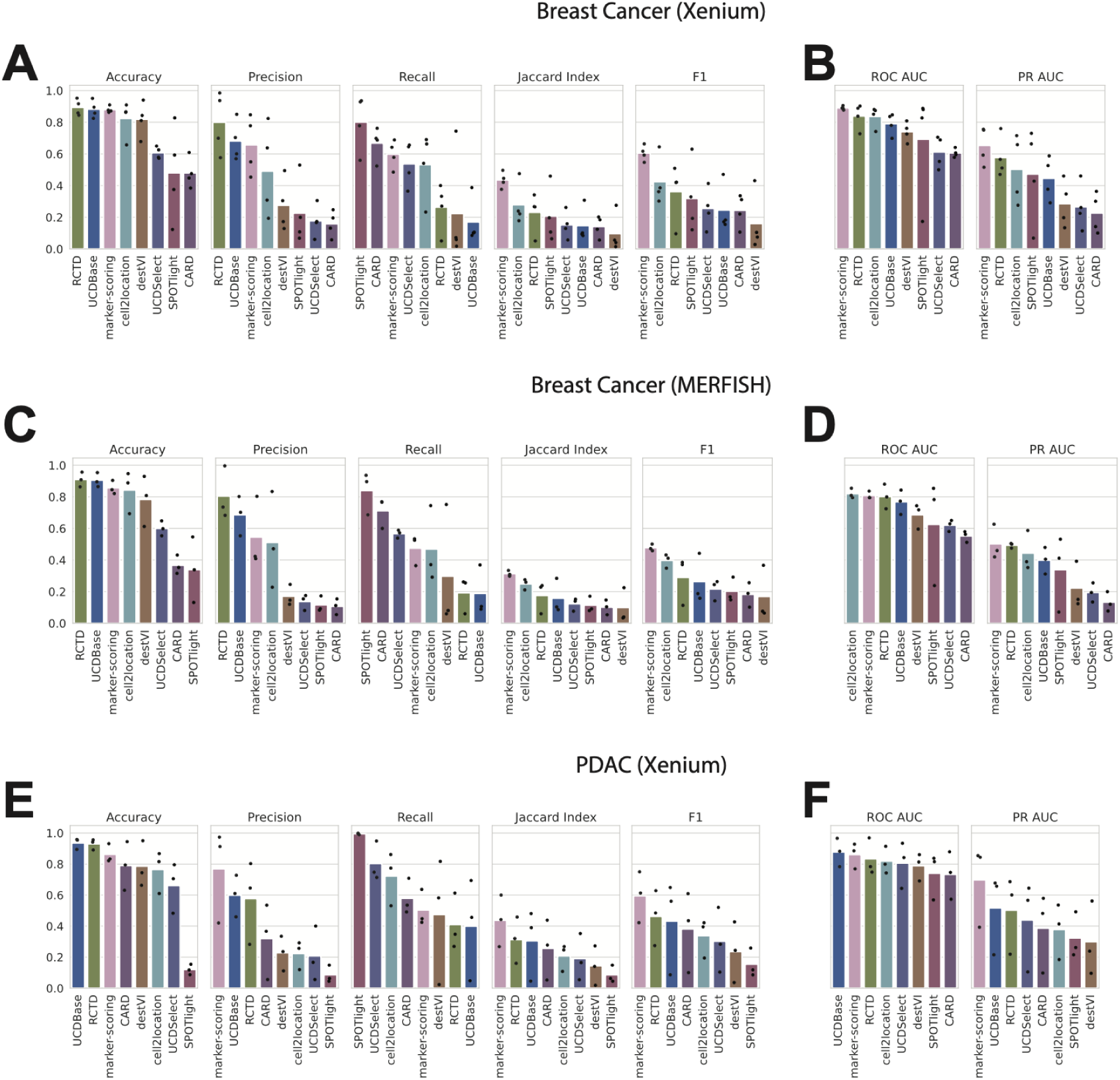
Benchmarking performance for rare cell types using binary classification metrics. A. Accuracy, precision, recall, Jaccard index, and F1 score for rare cell types across deconvolution methods in the Xenium breast cancer simulation. Predicted cell type proportions were binarized into presence or absence using a 5% threshold and compared to ground truth presence/absence for rare cell types. Bar heights represent the mean across cell types; individual values are overlaid as points. B. ROC AUC and PR AUC for rare cell types across deconvolution methods in the Xenium breast cancer simulation. Continuous predicted proportions were compared to binary ground truth. Bars represent mean values; individual values are overlaid. C. Same as panel A, but for the MERFISH breast cancer simulation. D. Same as panel B, but for the MERFISH breast cancer simulation. E. Same as panel A, but for the Xenium PDAC simulation. F. Same as panel B, but for the Xenium PDAC simulation.

**Figure S6.**
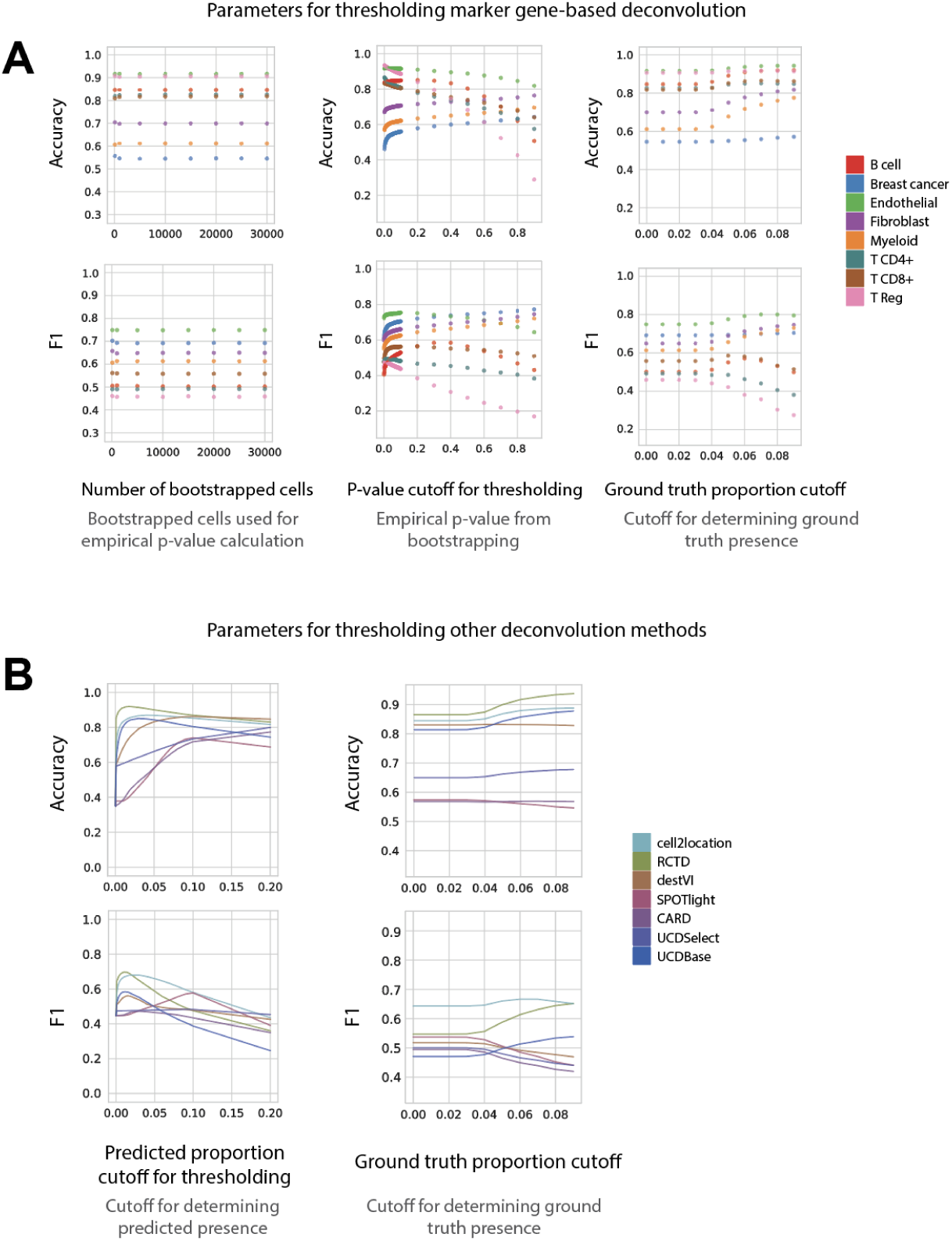
Parameter evaluation for marker gene scoring and deconvolution methods in predicting cell type presence. A. Scatterplots showing the impact of parameter variation on marker gene scoring performance. Accuracy (top row) and F1 score (bottom row) are plotted across a range of parameter values for marker gene-based presence/absence predictions, compared to binarized ground truth. Points are colored by cell type. The x-axis indicates the tested parameter values, and the y-axis shows the metric score. Evaluations were performed using the MERFISH breast cancer simulated dataset. B. Line plots showing the impact of parameter variation on deconvolution method performance. Accuracy (top row) and F1 score (bottom row) are plotted across a range of parameter values for presence/absence predictions, comparing deconvolution method outputs to ground truth. Each line represents the mean metric across cell types for a given method. The x-axis denotes the tested parameter values, and the y-axis shows the metric score. Evaluations were performed on the MERFISH breast cancer simulated dataset.

**Figure S7.**
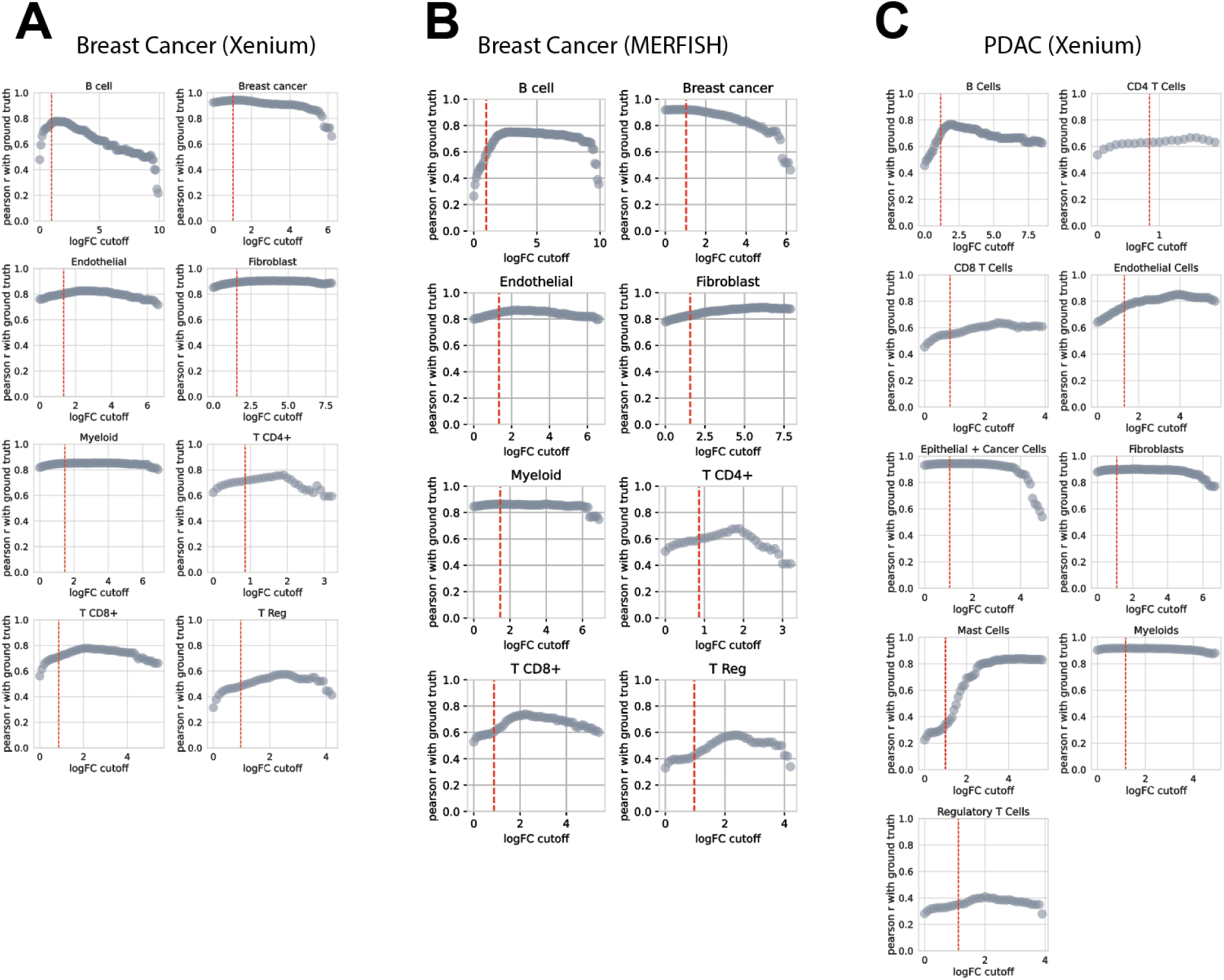
Effect of log fold-change cutoff on marker gene selection and deconvolution performance. A. Scatterplots showing the effect of logFC cutoffs used to define cell type-specific marker genes on deconvolution performance using marker gene scoring. The y-axis shows Pearson correlation between enrichment scores and ground truth cell type proportions. The red vertical line marks the 85th percentile (top 15%) of logFC values. Results shown for the Xenium breast cancer simulation. B. Same as panel A, but for the MERFISH breast cancer simulation. C. Same as panel A, but for the Xenium PDAC simulation.

**Figure S8.**
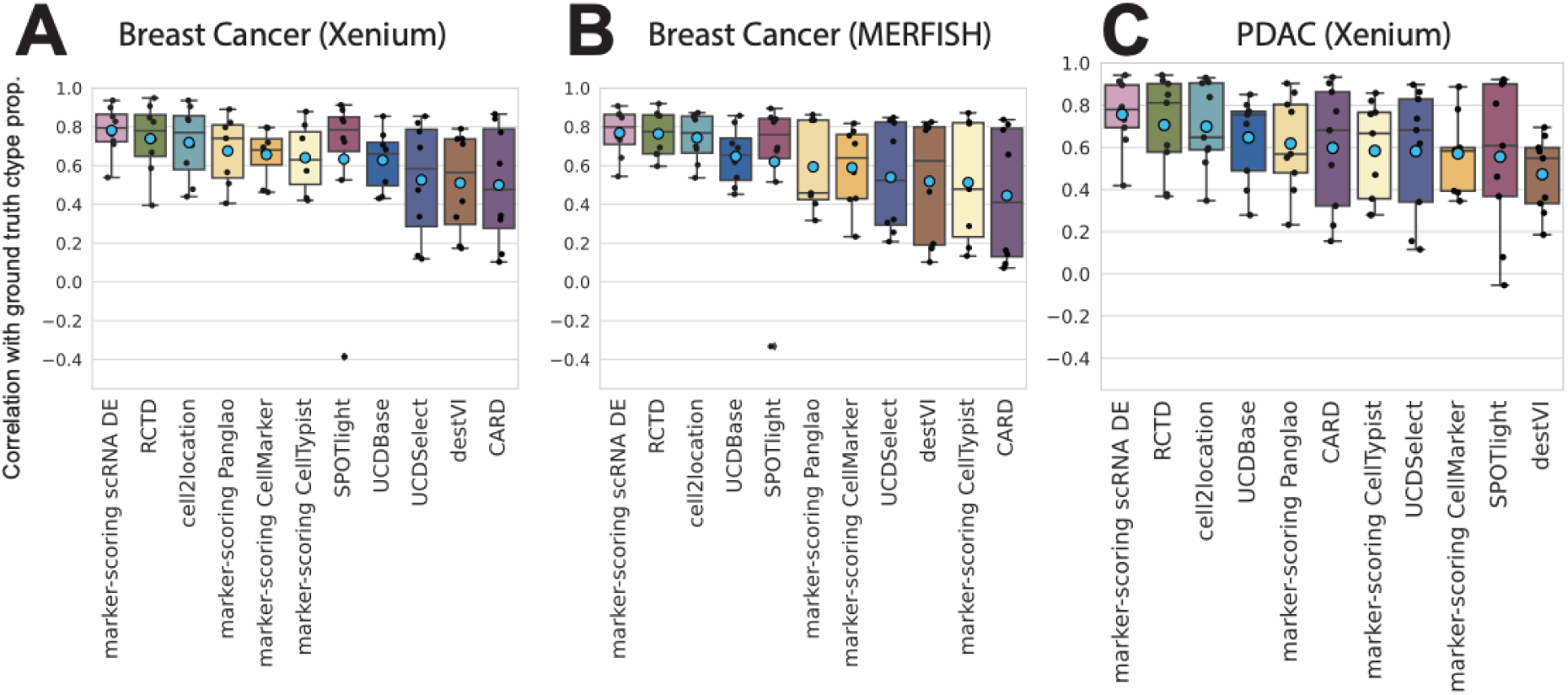
Benchmarking by marker gene source. A. Boxplots showing Pearson correlation between predicted scores from deconvolution methods and ground truth cell type proportions for the Xenium breast cancer simulation. Each point represents an individual cell type (stripplot overlay). For marker gene scoring, the x-axis denotes the source of marker gene sets. Blue diamond: mean; center line: median; box: interquartile range (IQR); whiskers: 1.5x IQR. B. Same as panel A, but for the MERFISH breast cancer simulation. C. Same as panel A, but for the Xenium PDAC simulation.

**Figure S9.**
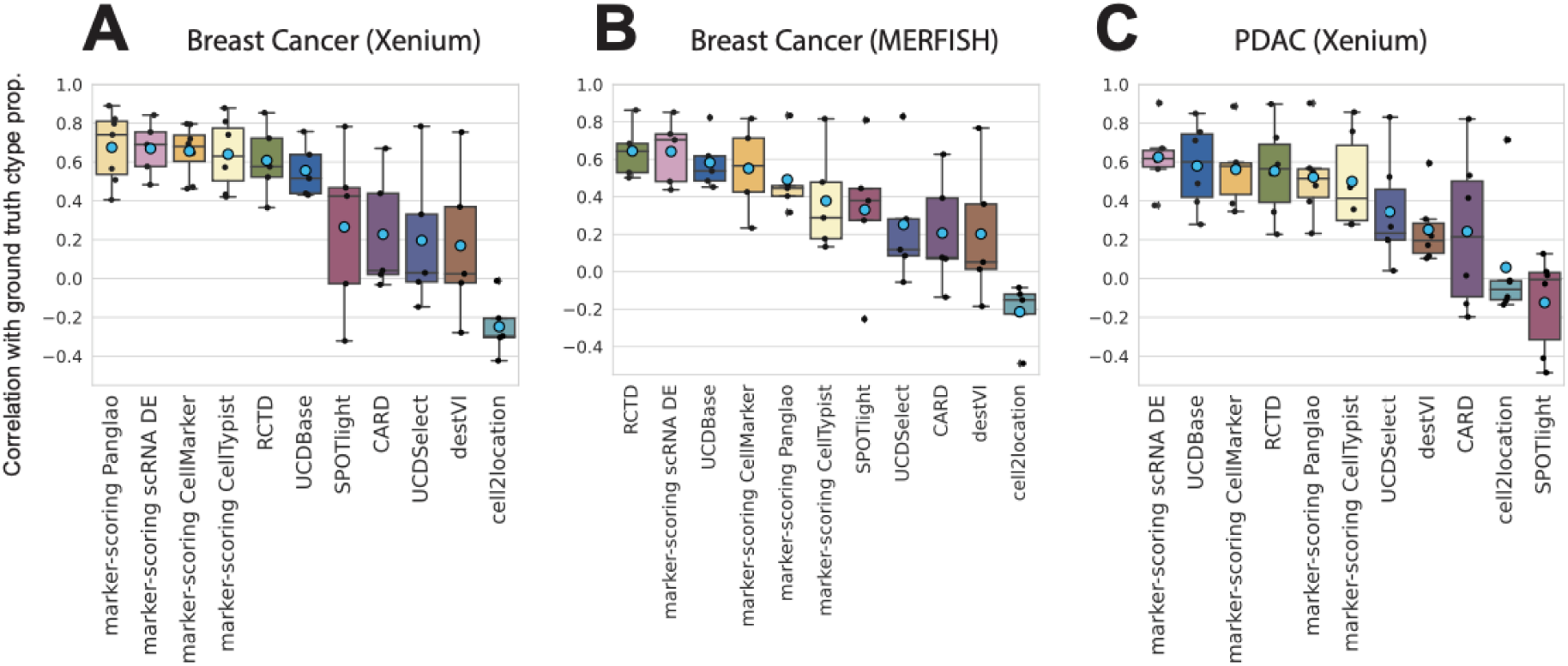
Benchmarking performance on immune-restricted references. A. Boxplots showing Pearson correlation between predicted cell type scores and ground truth proportions for the Xenium breast cancer simulation, when restricting the scRNA-seq reference to immune cell types only. Ground truth proportions were calculated using all cell types, while marker genes were derived from differential expression between immune cells in the reference. Individual cell types are overlaid as a stripplot. Blue diamond: mean; center line: median; box: interquartile range (IQR); whiskers: 1.5x IQR. For marker gene scoring, x-axis labels indicate the marker gene source. B. Same as panel A, but for the MERFISH breast cancer simulation. C. Same as panel A, but for the Xenium PDAC simulation.

**Figure S10.**
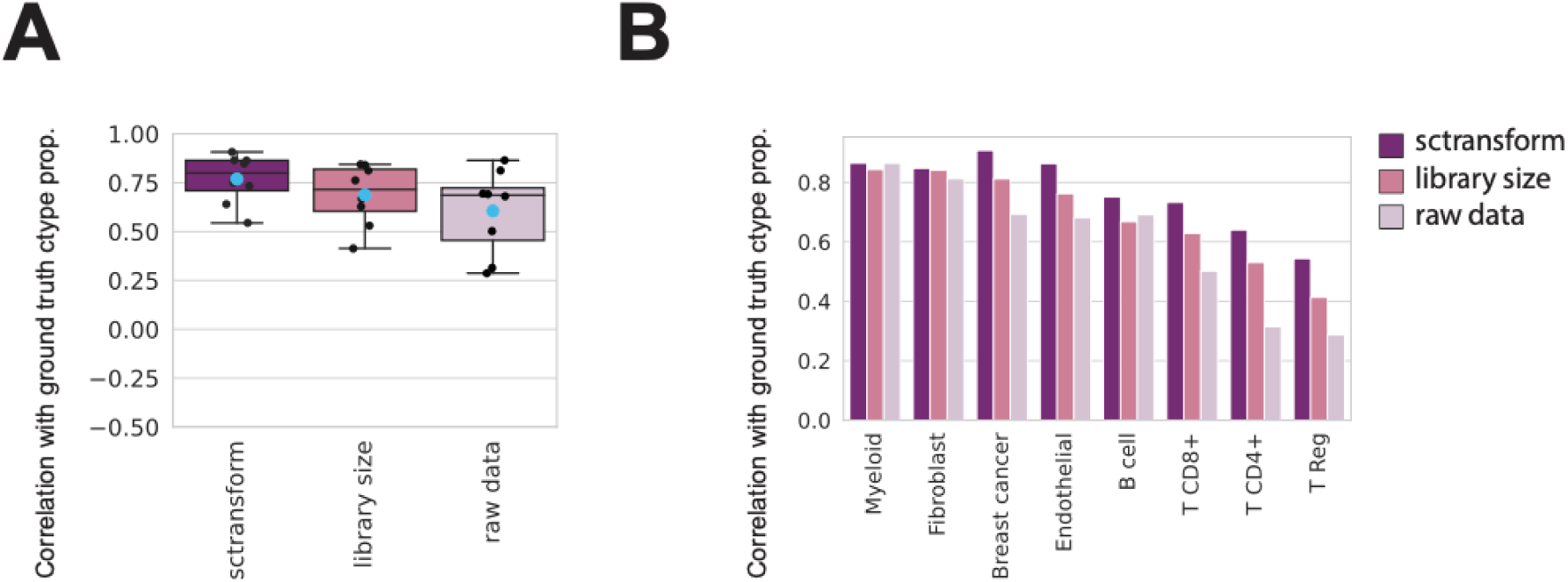
Impact of normalization method on marker gene scoring performance in Visium-like simulations. A. Boxplot showing the Pearson correlation between enrichment scores from bin-scoring and ground truth cell type proportions across normalization strategies in the MERFISH breast cancer simulation. Each point represents a single cell type. Blue diamond: mean; center line: median; box: interquartile range (IQR); whiskers: 1.5× IQR. B. Barplot showing the same Pearson correlation values as in panel A, grouped by cell type and colored by normalization method.

**Figure S11.**
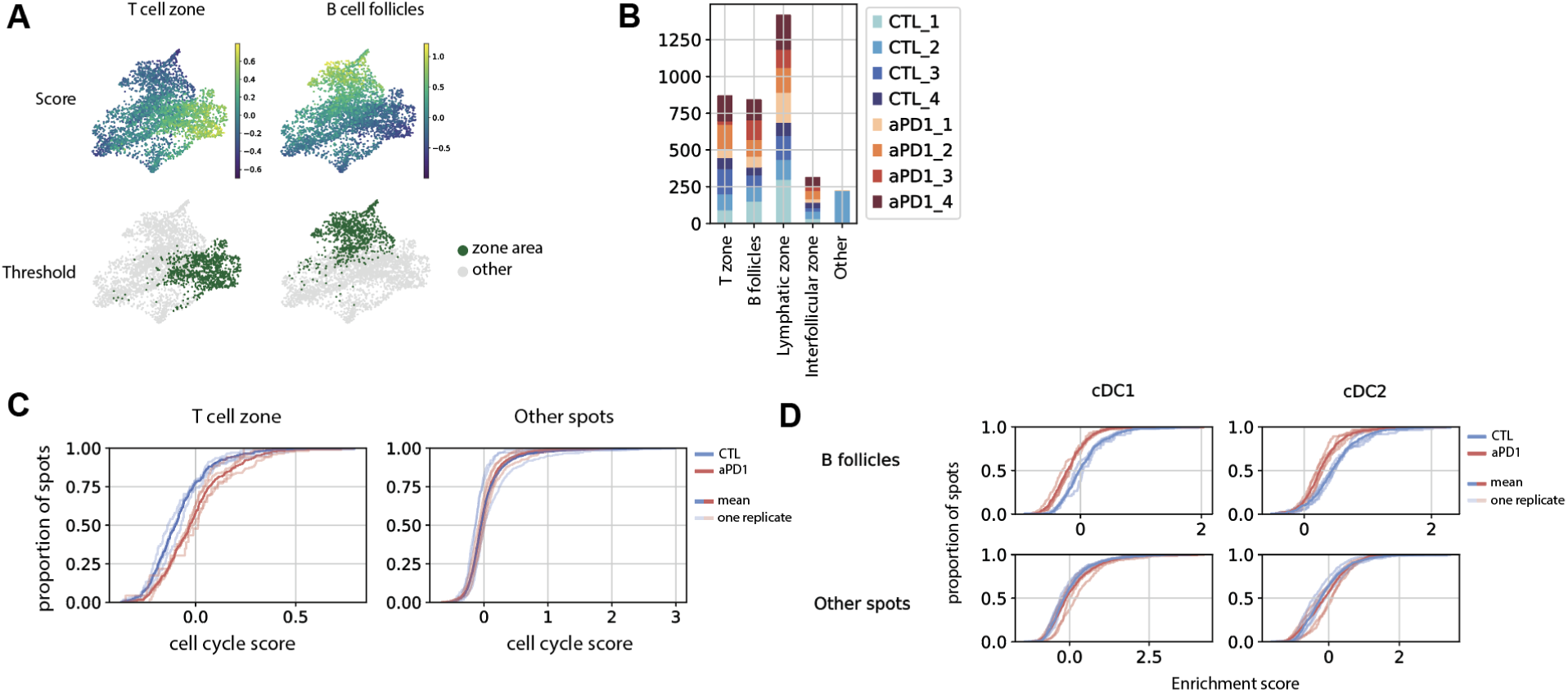
Tumor-draining lymph node (tdLN) data analysis. A. UMAP plots showing enrichment scores and binarized annotations for the T cell zone and B cell follicles across lymph node sections. B. Barplots showing the composition of annotated zones for each tdLN section. C. Cumulative distribution function (CDF) plots of the cell cycle scores in the T cell zone (left) and in all other regions (right). Each semi-transparent curve represents one biological replicate (pair of sections from the same lymph node), colored by treatment condition; solid curves represent all spots per condition. D. CDF plots for dendritic cell (DC) subtype scores (conventional dendritic cell type 1, cDC1; conventional dendritic cell type 2, cDC2) in the B cell zone (top) and other regions (bottom), with the same format of the plot as in panel C.

**Figure S12.**
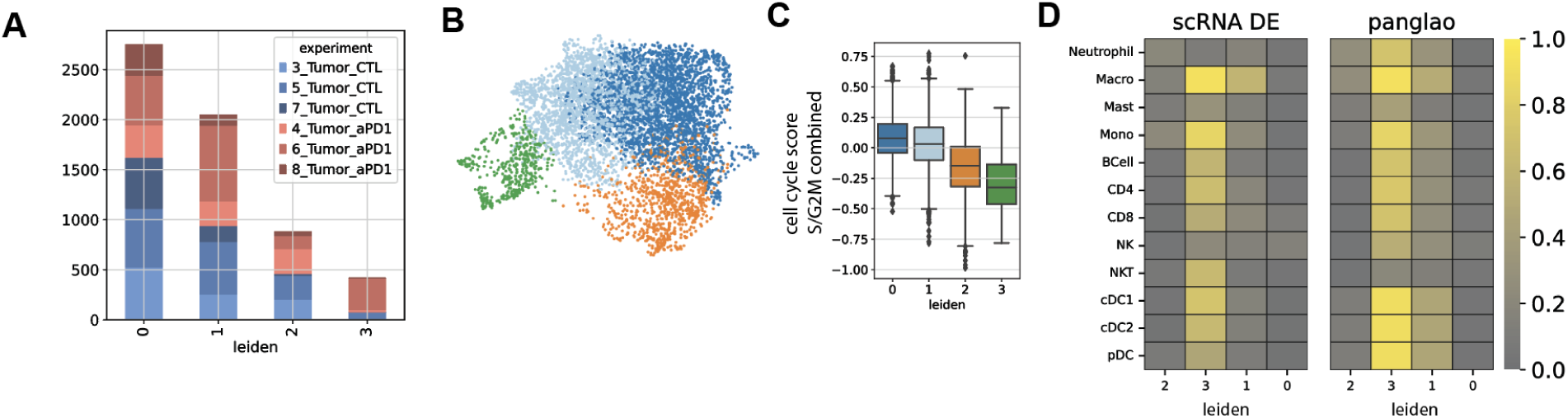
Tumor cluster overview and cell type composition. A. Barplot showing the number of spatial transcriptomic spots from each tumor sample assigned to each transcriptomic cluster. Clusters were defined using only transcriptomic data, without using spatial information. B. UMAP visualization of tumor Visium spots based on gene expression, with transcriptomic cluster annotations. C. Boxplot of per-spot cell cycle scores across tumor clusters. D. Proportion of spots in each cluster predicted to contain various cell types using marker gene signatures derived from scRNA-seq differential expression (left) or PanglaoDB (right).

**Figure S13.**
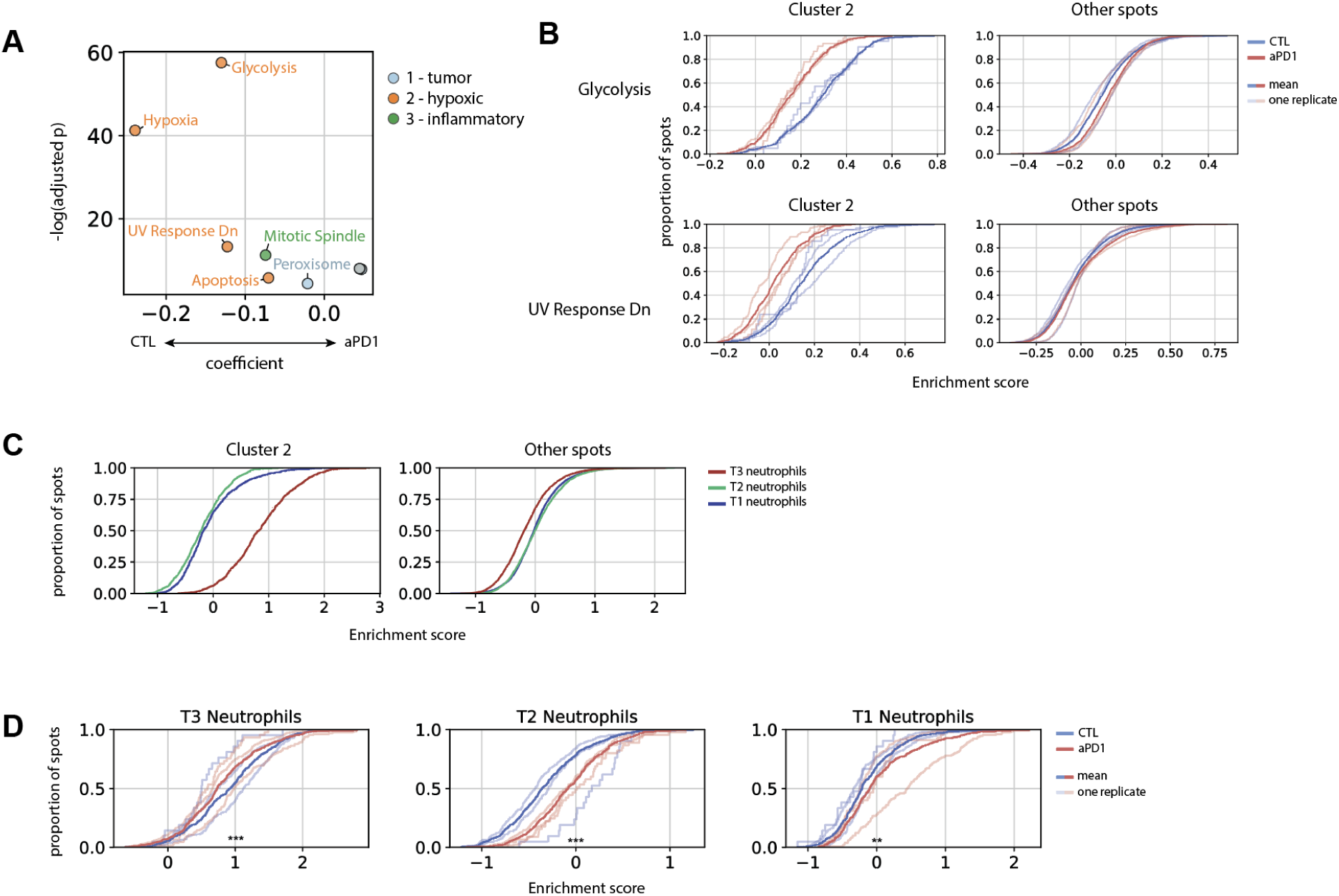
Tumor pathway and hypoxia analysis. A. Scatterplot of all significantly differential pathway scores between treatment conditions across tumor clusters. X-axis: condition coefficient from generalized linear model (GLM); Y-axis: –log10(p-value). Each point represents a pathway, colored by the tumor cluster in which it is differentially enriched. B. CDF plots for pathway scores in cluster 2 (left) and all other tumor spots (right). Each semi-transparent curve represents data from one mouse, colored by condition; solid lines show the distribution across all spots per condition. C. CDF plots of neutrophil subtype scores in cluster 2 (left) and all other spots (right). D. CDF plots for all neutrophil subtype scores in cluster 2, colored by treatment condition. Each semi-transparent curve shows a single mouse; solid lines represent condition-level distributions. Asterisks indicate statistical significance from two-sided t-test comparing the conditions: p < 0.05 (*), p < 0.01 (**), p < 0.001 (***).

